# Pleiotropy-robust Mendelian Randomization

**DOI:** 10.1101/072603

**Authors:** Hans van Kippersluis, Cornelius A Rietveld

## Abstract

**Background:** The potential of Mendelian Randomization studies is rapidly expanding due to (i) the growing power of GWAS meta-analyses to detect genetic variants associated with several exposures, and (ii) the increasing availability of these genetic variants in large-scale surveys. However, without a proper biological understanding of the pleiotropic working of genetic variants, a fundamental assumption of Mendelian Randomization (the exclusion restriction) can always be contested.

**Methods:** We build upon and synthesize recent advances in the econometric literature on instrumental variables (IV) estimation that test and relax the exclusion restriction. Our Pleiotropy-robust Mendelian Randomization (PRMR) method first estimates the degree of pleiotropy, and in turn corrects for it. If a sample exists for which the genetic variants do not affect the exposure, and pleiotropic effects are homogenous, PRMR obtains unbiased estimates of causal effects in case of pleiotropy.

**Results:** Simulations show that existing MR methods produce biased estimators for realistic forms of pleiotropy. Under the aforementioned assumptions, PRMR produces unbiased estimators. We illustrate the practical use of PRMR by estimating the causal effect of (i) cigarettes smoked per day on Body Mass Index (BMI); (ii) prostate cancer on self-reported health, and (iii) educational attainment on BMI in the UK Biobank data.

**Conclusions:** PRMR allows for instrumental variables that violate the exclusion restriction due to pleiotropy, and corrects for pleiotropy in the estimation of the causal effect. If the degree of pleiotropy is unknown, PRMR can still be used as a sensitivity analysis.

**Key messages:**

- If genetic variants have pleiotropic effects, causal estimates of Mendelian Randomization studies will be biased.
- Pleiotropy-robust Mendelian Randomization (PRMR) produces unbiased causal estimates in case (i) a subsample can be identified for which the genetic variants do not affect the exposure, and (ii) pleiotropic effects are homogenous.
- If such a subsample does not exist, PRMR can still routinely be reported as a sensitivity analysis in any MR analysis.
- If pleiotropic effects are not homogenous, PRMR can be used as an informal test to gauge the exclusion restriction.

## Introduction

Establishing causal effects of treatments or exposures on behavioral and disease outcomes is of great public health importance.^1^ The practice of “Mendelian Randomization” (MR) uses genetic variants as instrumental variables (IV) for a certain modifiable exposure in order to estimate the causal effect of that exposure on a certain disease or other outcome.^2,3^ This method has the potential to overcome the traditional biases due to confounding and reverse causality that plague observational studies.^4^ The past decade has seen increasing interest in MR, ^4,5,6, 7^ and its potential is rapidly developing through the increasing number of Genome-Wide Association Studies (GWAS) that find robust associations between genetic variants and exposures of interest.^8,9^

The assumptions of MR have been discussed extensively^2,10,11,12,13^, and the advantages and disadvantages of MR are heavily debated.^14,15^ Recently, there has been much progress in dealing with the disadvantages of MR, yet the critical assumption remains the exclusion restriction: the proposed IV (genetic variant) should not directly affect the outcome.^14,15^ While this assumption can be contested in any IV approach, the assumption is even more critical in the context of MR, since the biological working of genes is usually poorly understood.^13,16^

In particular, more and more studies show how the same genetic variant affects multiple outcomes through different biological pathways, a phenomenon known as biological (or horizontal) pleiotropy^17^, and this violates the exclusion restriction. In contrast, mediated (or vertical) pleiotropy, in which a genetic variant is associated with a certain phenotype and this phenotype is causal for a second phenotype, is not problematic.^4^ Therefore, we will only focus on biological pleiotropy, for brevity “pleiotropy” from here.

In response to the possible problem of pleiotropy, a couple of MR approaches have been proposed. Davey Smith and Hemani suggest using multiple genetic variants consecutively as IV, and argue that it is increasingly unlikely that 2, 3 or 4 different genetic variants produce the same estimate of the causal effect.^4^ While an appealing argument, this approach still hinges critically on the assumption that at least some of the genetic variants do not violate the exclusion restriction. Moreover, this informal test cannot discriminate between violations of the exclusion restriction and heterogeneous causal effects.^18^ Kang et al. propose a method that produces valid estimates in case at least 50% of the combined instrument strength across all variants originates from variants that satisfy the exclusion restriction.^19^ While helpful, it is generally not possible to distinguish the valid IVs from the invalid IVs in case the estimates based on different sets of IVs diverge.^20^ Moreover, this approach, like the weighted median estimator^21^, still requires some of the genetic variants to satisfy the exclusion restriction.

Bowden et al. propose to apply an Egger regression to Mendelian Randomization estimates.^20^ The idea is that a violation of the exclusion restriction leads to a bias of the MR estimate that is inversely proportional to the first-stage coefficient of the IV on the exposure. Under the assumption that, across all genetic variants, the effect of the IV on the outcome and the effect of the IV on the exposure are uncorrelated (“InSIDE assumption”), this implies that IVs with a stronger effect on the exposure should give less-biased MR estimates. A regression of the MR estimates on the first stage coefficients including an intercept then provides a consistent estimate of the causal effect. As acknowledged by the authors, this InSIDE assumption cannot be tested and may not hold if the genetic variants used as IVs are correlated with confounders of the association between exposure and outcome.

In this paper we build upon, and combine, two recent advances in the econometric literature on IV estimation that test and relax the exclusion restriction. We introduce the “plausibly exogenous” method^22^ in MR research to account for pleiotropy. Two studies have previously noticed the possibility of applying this method as a sensitivity analysis in the context of MR, ^12,23^ but both papers do not provide any guidance on how to choose the essential input parameters. Our innovation is that we combine this method with another stream of econometric research that designs auxiliary regressions to test for violations of the exclusion restriction.^24,25,26^ The intuition is that in a subsample for which the first stage (that is, the effect of the IV on the exposure) is zero, the reduced form (that is, the effect of the IV on the outcome) should be zero too in case the exclusion restriction is satisfied. While traditionally used merely as a test of the exclusion restriction, there is no earlier notion that the reduced-form estimate obtained in this subsample is exactly the input required for the “plausibly exogenous” method. We term the synthesis of these techniques Pleiotropy-robust Mendelian Randomization (PRMR).

Simulation results show that if a sample exists for which the first stage is zero, and the pleiotropic effects are homogenous, it is possible to obtain unbiased estimates of causal effects using PRMR, even when all genetic instruments violate the exclusion restriction. We empirically illustrate our method by estimating the effect of (i) the number of cigarettes smoked per day on body mass index (BMI), (ii) prostate cancer on subjective health evaluations, and (iii) educational attainment on BMI.

## Methods

### Mendelian Randomization

In the general case, one is interested in the causal effect *β* of a certain exposure *X* on an outcome *Y*. The idea of MR is that there is a vector of genetic variants *G* (usually single nucleotide polymorphisms (SNPs), an allele score, or polygenic score) that is known to be correlated with the exposure *X*, but is assumed to be uncorrelated with other (unobserved) determinants of the outcome *Y*. Consider the equations (we follow the notation of Bowden et al.^20^ in matrix notation here)

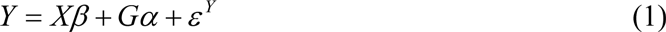

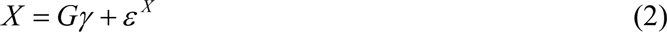

where *ε^Y^* and *ε^X^* are composite error terms including unobserved confounders *U*. The assumptions of MR are (see Figure 1 for a graphical exposition):^12,13,16^

1. Relevance: The genetic variants *G* have an effect on the exposure *X*: *γ* ≠ 0.
2. Independence: The genetic variants *G* are uncorrelated with any confounders of the exposure-outcome relationship.
3. Exclusion restriction: The genetic variants *G* affect the outcome *Y* only through the exposure *X*: *α* = 0.

**Figure 1.**
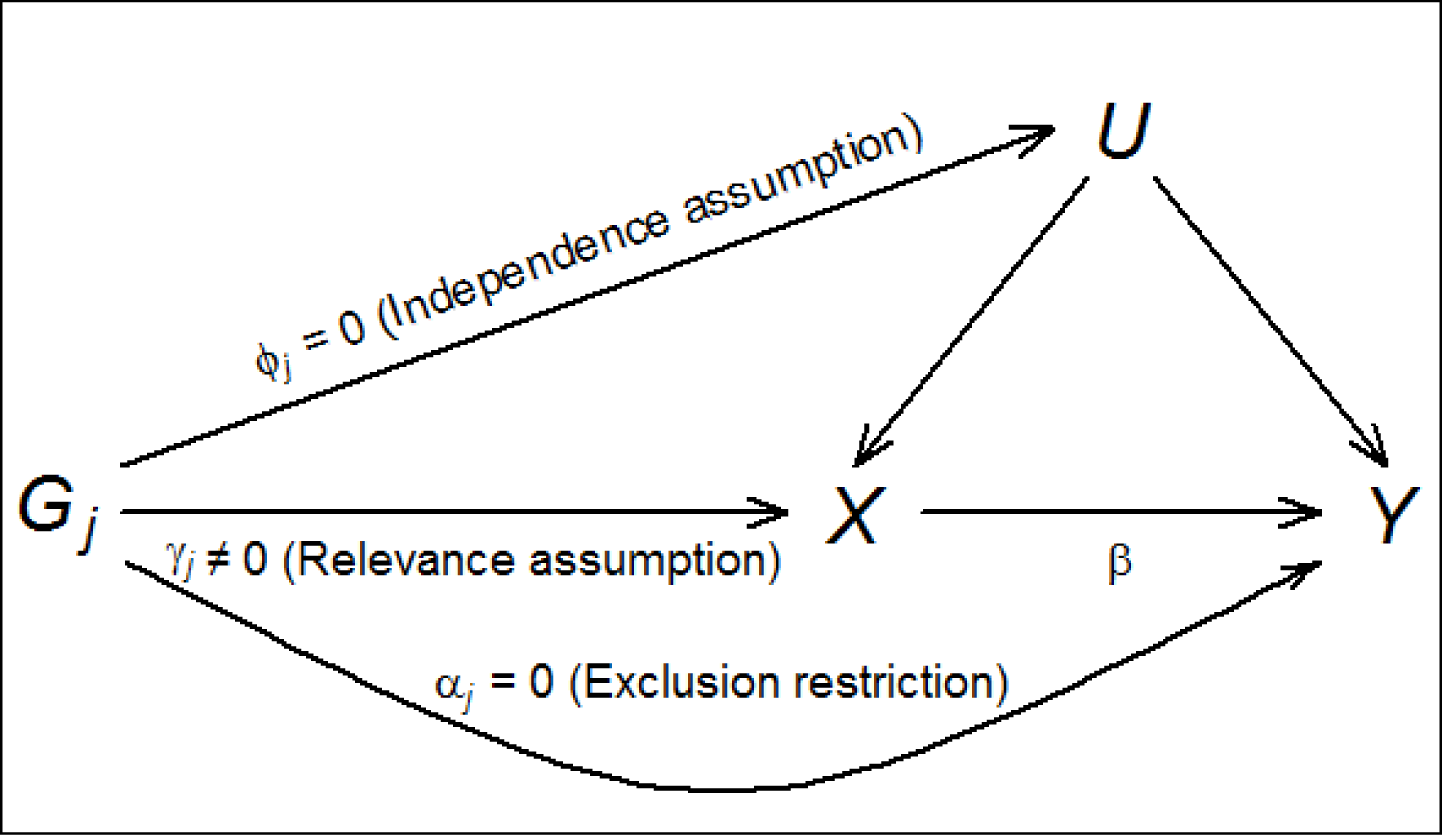
Illustrative diagram showing the standard MR model with its assumptions. The genetic effect of instrumental variable *G_j_* on exposure *X* is *γ_j_*; the genetic effect on unobserved confounder *U* is *ϕ_j_*; the direct genetic effect on the outcome *Y* is *α_j_*, and the causal effect of the exposure *X* on the outcome *Y* is *β*.

The use of genetic variants as IV has at least two very attractive properties. First, publicly available GWAS results make it relatively straightforward to select genetic variants *G* for which *γ* ≠ 0, i.e. genetic variants that are robustly associated with the exposure of interest. Second, given that genetic variants are randomly distributed at conception, conditional on population stratification variables or family-specific effects, it is usually plausible that the independence assumption holds.^27^

The exclusion restriction is widely acknowledged as the most problematic assumption of MR.^14, 15^ In particular, the existence of pleiotropy would lead to a violation of the exclusion restriction, and *α* ≠ 0 in equation (1). In traditional MR it is assumed however that *α* is equal to 0, which leads to biased estimates of *β*, the causal effect of interest, in case pleiotropy is present. Moreover, this bias gets amplified by the typically low explanatory power of the genetic variants for the exposure.^28^

### Pleiotropy-robust Mendelian Randomization (PRMR)

In the “plausibly exogenous” method^22^ the assumption that *α* = 0 is relaxed, and replaced by a user specified assumption on the plausible value, range or distribution of *α*. When the prior on *α* follows a Normal distribution with mean *μ*_*α*_ and variance Ω_*a*_, and the uncertainty about *α* reduces with the sample size (i.e., “local-to-zero”), then the estimate of the causal effect *β* in equation (1) is given by:

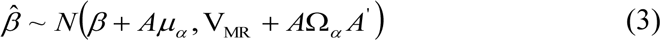

Where *N()* indicates the Normal distribution, *A* = (*X*′*G*(*G*′*G*)^-1^*G*′*X*)^-1^(*X*′*G*), and *β* and *V_MR_* are the traditional MR point estimate and variance-covariance matrix, respectively. The plausibly exogenous method on itself, however, provides no guidance on the value or distribution of *α*.

We can, however, estimate the value of *α* if there is a population subgroup for which the first stage is known to be zero. A recent stream of econometric research emphasizes the identification of these subgroups to test the exclusion restriction.^24,25,26^ An early example is Altonji et al. (2005)^24^, who investigate the validity of the instrument ‘being Catholic’ to study the effect of attending a Catholic high school on a wide variety of outcomes. They identify a subsample of public eighth graders among which practically nobody subsequently attends a Catholic high school. Hence, among this subsample the first stage is zero, and any association between the IV (being Catholic) and the outcome reflects a direct effect, indicating a violation of the exclusion restriction.

While this strategy is so far exclusively used as a test of the exclusion restriction, our observation is that the result of this auxiliary regression is exactly the required input for the “plausibly exogenous” method. Consider the reduced form equation that is obtained by substituting (2) into (1):

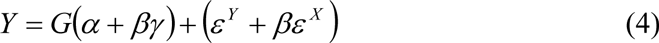

In a subsample for which the first stage is zero (*γ* = 0), the reduced form coefficient of the genetic variant is an estimator for *α*.

Practically, we suggest first estimating the reduced-form (4) in a sample for which *γ* = 0, to obtain estimates 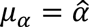 and Ω_*α*_ equal to the squared standard error of 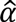. Thereafter, plug in *μ*_*α*_ and Ω_*α*_ into the plausibly exogenous equation (3), to obtain estimates of the causal effect of interest *β*. The estimator is easy to obtain in standard software. For example, the user-written command “plausexog” is readily available in STATA.^29^ While the plausibly exogenous method has been originally developed for continuous outcomes, it is also valid for binary outcomes in as far one is willing to estimate linear probability models for binary outcomes.

### Simulation

We present a simulation study to illustrate the performance of regular MR through Two-Stage Least Squares (2SLS), the Inverse Variance Weighting (IVW) method^30^, MR-Egger regression, and PRMR. Following Bowden et al. (2015)^20^, we consider the following four scenarios with varying violations of the MR assumptions (see the appendix for more details):

1. No pleiotropy
2. Balanced pleiotropy, InSIDE satisfied (*α* parameters take positive and negative values, but are independent of the first stage parameters *γ*)
3. Directional pleiotropy, InSIDE satisfied (*α* parameters take only positive values, but are independent of the first stage parameters *γ*)
4. Directional pleiotropy, InSIDE not satisfied (*α* parameters take only positive values and are correlated with the first stage parameters *γ*)

As in Bowden et al. (2015) we assume that all of the SNPs violate the exclusion restriction in scenarios 2-4, and so we do not consider methods that require at least 50% of the SNPs to be valid. In all scenarios, we use a sample size of 1,000; 10,000 simulation runs; 25 genetic variants with minor allele frequency 0.30; and a causal effect *β* of 0.00 and 0.05.

**Table 1.** Performance of Two-stage-least-squares (2SLS), Inverse-variance weighting (IVW), MR-Egger regression and PRMR in simulation study for one-sample Mendelian randomization with a null (*β* = 0.00) and a positive (*β* = 0.05) causal effect.

| Scenario | Mean estimate |  |  |  |
| --- | --- | --- | --- | --- |
|  | 2SLS | IVW | MR-Egger | PRMR |
| $\beta = 0.00$ | | | | |
| 1: No pleiotropy, InSIDE satisfied | 0.00 | 0.00 | 0.01 | 0.00 |
| 2: Balanced pleiotropy, InSIDE satisfied | 0.00 | 0.00 | 0.01 | 0.00 |
| 3: Directional pleiotropy, InSIDE satisfied | 0.04 | 0.04 | 0.01 | 0.00 |
| 4: Directional pleiotropy, InSIDE violated | 0.13 | 0.13 | 0.04 | 0.00 |
| $\beta = 0.05$ | | | | |
| 1: No pleiotropy, InSIDE satisfied | 0.05 | 0.05 | 0.06 | 0.05 |
| 2: Balanced pleiotropy, InSIDE satisfied | 0.05 | 0.05 | 0.06 | 0.05 |
| 3: Directional pleiotropy, InSIDE satisfied | 0.09 | 0.09 | 0.06 | 0.05 |
| 4: Directional pleiotropy, InSIDE violated | 0.18 | 0.18 | 0.09 | 0.05 |

The simulation results show that in a one-sample setting 2SLS and IVW produce very similar results, and that in case of no pleiotropy and balanced pleiotropy all methods give average estimates of *β* (almost) equal to the true value. With directional pleiotropy (both when InSIDE is satisfied and violated), the average estimates are biased away from the true value for 2SLS and IVW. MR-Egger performs slightly better, but still produces biased estimates. Not surprisingly, when the nature of pleiotropy is known, PRMR gives unbiased estimates. We acknowledge however that both deviations from homogenous pleiotropic effects, and non-zero first-stage coefficients among the subsample for which the first stage should be zero, would produce biased estimates for *α*, and in turn the causal effect *β*.

## Examples

To illustrate our approach, we exploit genetic data from the interim release of the UK Biobank^31^ to study (i) the effect of the number of cigarettes smoked per day (CPD) on BMI; (ii) the effect of prostate cancer on subjective health evaluations; and (iii) the effect of educational attainment on BMI. Following recommendations from the genotyping center, we restrict the analyses in the UK Biobank to 112,338 (52,53% female) conventionally unrelated individuals with “White British” ancestry.^32^ We use the first 15 principal components (PCs) of the genetic relatedness matrix as provided by UK Biobank to further control for population stratification.^32,33^ All individuals are born between 1934 and 1970. For individuals with missing information on a specific SNP, we impute the mean genotype from the sample. For all measures, we use reported values from the first interview round.

### The effect of Cigarettes per Day on BMI

The relationship between smoking and BMI has received considerable attention in the literature^34,35,36,37,38^, and from these studies Wehby et al. (2012)^37^ used MR to assess the causal effect of CPD on BMI. In existing GWAS, the SNPs that are robustly associated with smoking measures can mostly be traced back to nicotine dependence.^39^ This provides a context to apply PRMR: after all, the development of nicotine dependence requires initiating smoking in the first place. Hence, the group of never smokers provides a subsample among which the SNPs do not have an effect on CPD, and we can use this subsample to estimate the direct effect of the SNPs on the outcome measure BMI.

As instrument we use standardized values of rs12914385, the SNP with the strongest statistical association with CPD from the only locus found to be associated with CPD in the GWAS from the Tobacco and Genetic Consortium.^40^ CPD is measured as number of cigarettes currently smoked per day, and BMI is measured in kg/m^2^. OLS results suggest a positive association between CPD and BMI (0.05, *p* = 1.90×10^-17^). The first stage regression of CPD on the SNP shows a strong positive association (*p* = 3.64×10^-16^, *F* = 66.47). Among never-smokers, the SNP has a very modest positive association with BMI (0.01, *p* = 0.44). In contrast, among current-smokers, there is a strong negative association with BMI (-0.16, *p* = 8.49×10^-5^). This provides evidence that the exclusion restriction is satisfied, and regular MR can be applied. The 2SLS results indicate a negative causal effect of CPD on BMI (-0.24, *p* = 3.50×10^-3^), suggesting an effect of CPD on BMI in the opposite direction of the OLS estimate.

### The effect of Prostate Cancer on Self-reported Health

Earlier studies have reported on the effect of (prostate) cancer on health outcomes^41,42,43^, yet none of these studies used Mendelian Randomization. Al Olama et al. (2014)^44^ find 12 autosomal SNPs to be related to prostate cancer at genome-wide significance level among individuals from European descent. Since prostate cancer naturally is only a risk factor among *males*, the first stage (that is, the effect of genetic variants on prostate cancer) among *females* is zero. As a result, the effect of a genetic variant associated with prostate cancer on a certain outcome among females may be used as a reasonable estimate for *α*.

UK Biobank contains a self-report about whether a person was told by a doctor to have prostate cancer (1) or not (0). Subjective health is measured on a four-point scale ranging from Excellent (4), via Good (3) and Fair (2), to Poor (1). We build an allele score for prostate cancer using the results from the GWAS on prostate cancer, ^44^ since the allele score has more power than using the individual SNPs.^45,46,47^ 10 out of the 12 SNPs are available in the genetic data of UK Biobank. The allele score has been standardized to have mean 0 and standard deviation 1.

An OLS regression of subjective health on prostate cancer among males reveals that prostate cancer is negatively associated with subjective health (-0.17, *p* = 9.17×10^-11^), and a regression of prostate cancer on the allele score shows that the allele score is positively associated with prostate cancer (*p* = 2.45×10^-3^, *F* = 9.21). A standard 2SLS regression with the allele score as instrument for prostate cancer produces a large negative effect of prostate cancer on subjective health (-1.26), but *p* = 0.54.

Using PRMR, we find that the allele score is negatively associated with subjective health among females (-0.009, *p* = 1.81×10^-3^), and this reduced form estimate is even larger in absolute terms than among males (-0.002, *p* = 0.53). This suggests that the exclusion restriction is violated and that the MR results are biased. When we plug in the reduced form estimate among females in the plausibly exogenous method we find that the effect of prostate cancer on subjective health is estimated to be positive (4.53) among males, but this estimate is implausibly large and surrounded by a large 95% confidence interval (-0.88 to 9.94, *p* = 0.10). Hence, we cannot reject a zero effect of prostate cancer on self-reported health.

### The effect of Educational Attainment on BMI

The education-health gradient is well-documented and one of the most robust findings in social science.^45, 49^ Several studies reported before on the causal effect of educational attainment on BMI, with mixed findings, and apart from one study investigating the reverse effect of BMI on educational attainment^50^, none of them used Mendelian randomization.^51,52,53,54,55,56^

Educational Attainment (EA) is constructed as years of education, similar as in the recent Educational Attainment GWAS (the UK Biobank data is used as out-of-sample replication sample in that study).^57^ For educational attainment, we build an allele score from the 74 SNPs identified found to be associated with EA.^57^ Of these 74 SNPs, 72 are available in UK Biobank.

An OLS regression of BMI on EA reveals that EA is negatively associated with subjective health (-0.11, *p* = 9.20×10^-308^), and a regression of EA on the allele score shows that the allele score is positively associated with EA ( *p* = 4.10×10^-122^, *F* = 552.12). A standard 2SLS regression with the allele score as instrument for EA provides a negative estimate (-0.39) for the causal effect of EA on BMI, with *p* = 9.63×10^-21^. However, it is likely that the exclusion restriction is violated, since the allele score for EA is, conditional on population stratification controls, gender, and birth year, positively associated with birth weight (0.01, *p* = 0.04), a known confounder of the relationship between EA and BMI.^58^

In contrast to the previous examples, here it is difficult to find a subsample for which the first stage is zero. Therefore we illustrate how PRMR can be used as a sensitivity analysis to determine how strong the violation of the exclusion restriction should be to render the causal effect *β* to be 0. Since one usually selects SNPs as IV for their association with the exposure rather than the outcome, it seems plausible that, in absolute value, the standardized first stage effect of the IV on the exposure, *γ*, is larger than the standardized direct effect of the IV on the outcome, *α*. Therefore, we define the proportion 0 ≤ *λ* ≤ 1, we set *μ*_*α*_ equal to 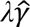, and the variance Ω_*α*_ equal to the squared standard error of 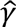. Subsequently, we apply equation (3) for different values of *λ* to find for which *λ* the corresponding *β* equals 0. When the first stage estimate has the opposite sign of the direct effect estimate, one needs to change the sign of 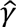 in this sensitivity analysis.

In our case, the first stage effect (allele score on EA) has plausibly the opposite sign of the direct effect (allele score on BMI) and hence we set *μ*_*α*_ equal to 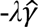. The resulting estimates for *β* are plotted in Figure 2 as a function of *λ*. The estimate in case of *λ* = 0 corresponds to the MR point estimate, but the 95% confidence interval is a little wider because 0 Ω_*α*_. Moving along the *x*-axis, the causal effect of EA on BMI is estimated to be 0 when *λ* = 0.41, and the 95% confidence interval includes 0 already when *λ* = 0.29. Hence, a relatively mild violation of the exclusion restriction – 29% of the first-stage effect – implies that we cannot reject a zero effect of EA on BMI, producing at best weak evidence that EA causally reduces BMI.

**Figure 2.**
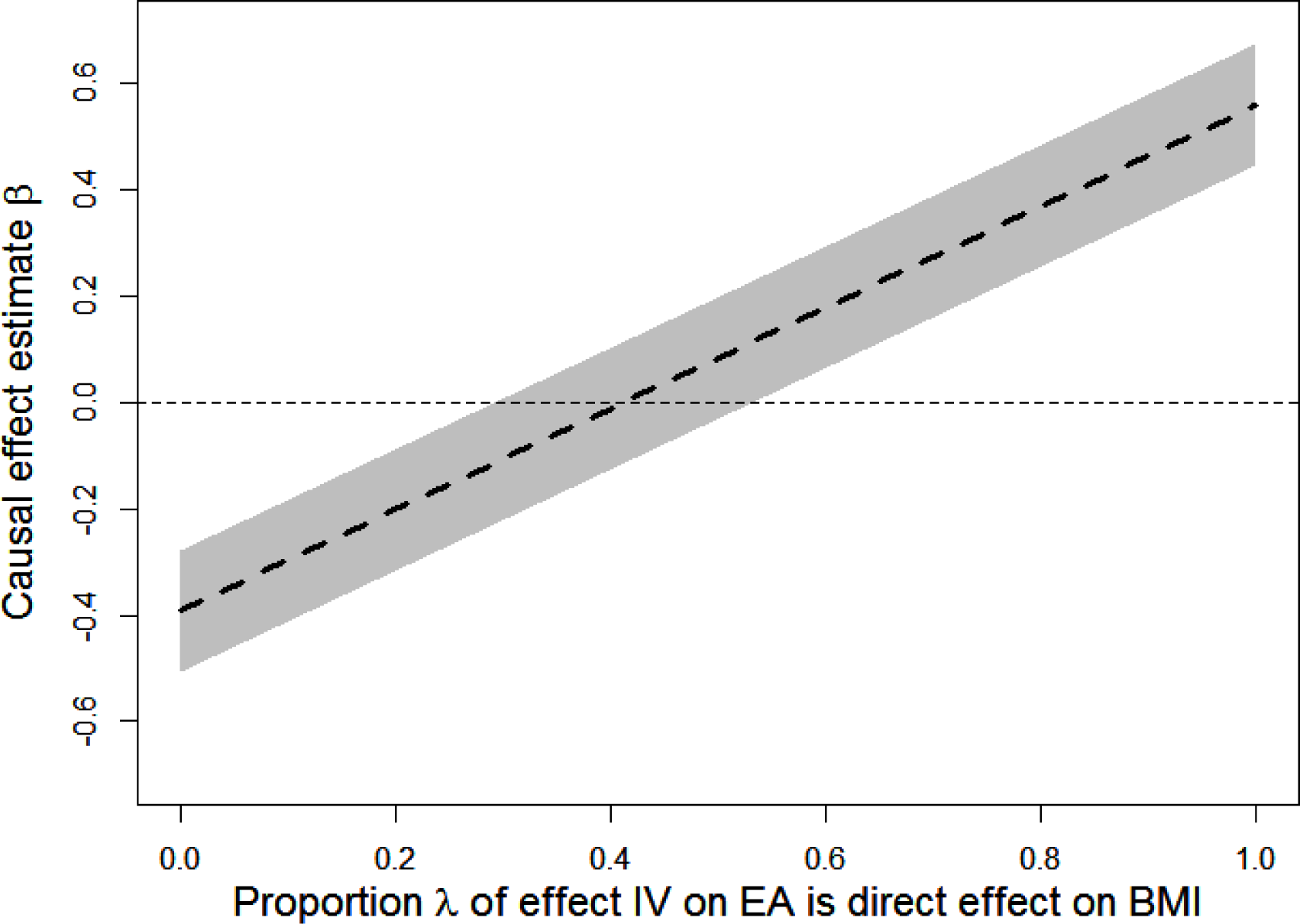
The causal effect of Educational Attainment on BMI, for varying values of λ (the percentage of the standardized effect of the IV on EA which is considered to be the direct effect of the IV on standardized BMI). The grey area represents the 95% Confidence Interval.

## Conclusion

The fact that the pleiotropic effects of genes are poorly understood makes it difficult to use genetic variants as credible instrumental variables in Mendelian Randomization. In this paper we argued that if (i) one can identify a subsample for which genetic variants do <u>not</u> have an effect on the exposure, and (ii) pleiotropic effects are homogenous, PRMR provides a way to deal with violations of the exclusion restriction due to pleiotropy, and opens the door for several applications in epidemiology and social science research.

A simulation study showed that PRMR clearly outperforms existing methods in case of violations of the exclusion restriction. We illustrated our PRMR approach by estimating the causal effect of cigarettes smoked per day on BMI, and the effect of prostate cancer on subjective health evaluations. In those two cases it is possible to identify subsamples where the effect of SNPs on the exposure are zero (never smokers and females, respectively), and this allows estimating, and if necessary correcting for, the pleiotropic effect.

We acknowledge that the two requirements for PRMR are not always satisfied. Yet, even if one is not willing to make the assumption of homogenous pleiotropic effects, one can still use the subsample without an effect of the genetic variants on the exposure as a useful test of the exclusion restriction. Moreover, if one cannot identify a subsample without a first stage, as we illustrated for the estimation of the effect of education on BMI, PRMR still allows for an informative sensitivity analysis that could routinely be applied in all MR analyses.

## Acknowledgments

This research has been conducted using the UK Biobank Resource. We thank Daniel Benjamin, Tom DiPrete, Patrick Turley, Peter Visscher, and Dinand Webbink for helpful comments.

